# Female behavioral cues control male courtship plasticity independent of female pheromones in African *Drosophila melanogaster*

**DOI:** 10.64898/2026.08.31.746996

**Authors:** Samuel Marston, Cadyn Mathis, Tamara Starling, Mariana Maya Galan, Andrea Agustin, Daniel A. Barbash, Dean M. Castillo

## Abstract

Sexual interactions involve the exchange of information—including olfactory, auditory, and visual cues—that has been shaped by sexual selection to maximize fitness. While these courtship signals and behaviors are often characterized as stereotypical and species-specific, male courtship is plastic and can depend on the population identity of the courted female. The ability of a male to adjust his courtship behavior based on female identity has significant implications for how selection operates on courtship, potentially driving behavioral divergence between populations and the eventual emergence of new species. A major challenge is identifying the cues that males use to adjust their courtship strategy. We used *Drosophila melanogaster* to investigate the mechanisms facilitating courtship plasticity, focusing on male song production. Male song is often considered the paramount behavior performed by males and is stimulated by female pheromones. Previous observations have focused on cosmopolitan (non-African) genotypes, whereas the singing rate and singing plasticity is highly variable in African populations. In genotypes from Zimbabwe, females have a different major pheromone compound (5,9-HD) compared to non-African females and show strong behavioral preference for Zimbabwe males. We created a Zimbabwe genotype lacking 5,9-HD by disrupting the gene *desat2* in the Zimbabwe Z53 strain. We tested the hypothesis that Zimbabwe male courtship plasticity depends on female pheromones by allowing males to court non-African, wild type Z53, and Z53 *desat2* null mutant females. We found that males treated Z53 *desat2* null females like wild type Z53 females, not using pheromone cues to determine singing and courtship plasticity. This suggests that other female behavioral feedback influences singing plasticity. This courtship plasticity could contribute to the asymmetrical reproductive isolation between these populations and could provide insight to the broad pattern of asymmetrical reproductive isolation across broad taxonomic groups.

## Introduction

During sexual interactions, males and females exchange signals and information that have been shaped by sexual selection to increase reproductive success and fitness (Mitoyen, et al. 2019; Wachtmeister 2001; Ritchie 2007; Shuster 2009). Information on mating status, receptivity, and species identity is communicated during these interactions, informing mate choice and mating outcomes (Perring, et al. 1993; Jennions and Petrie 1997; Rodriguez, et al. 2004). This information can be transmitted in multiple sensory modes including olfaction, such as pheromones, and auditory and visual behaviors (Uetz, et al. 2009; Higham and Hebets 2013; Oh and Shaw 2013). Together, these behaviors make up what is commonly referred to as courtship (Darwin 1872; Andersson 1994; Greenspan and Ferveur 2000). A main open question in evolutionary biology is how these signals have evolved such that different information stimulates courtship displays across populations and species.

Male courtship is often thought to be stereotypical, that is easily predicted and occurring in a relatively linear fashion (Ewing and Bennet-Clark 1968; Cobb and Jallon 1990; Kohatsu, et al. 2011; Yamamoto and Ishikawa 2013). Species designation can often be ascribed based on specific behaviors, courtship songs, or pheromone profiles (Kirkpatrick 1982; Crews and Moore 1986; Coyne and Orr 2004; Ritchie 2007; Hoskin and Higgie 2010). This contrasts with other social behaviors that are known to be context dependent and can change based on specific social environments, making these behaviors plastic (Bell, et al. 2009; Oliveira 2009; Froemke and Young 2021). While courtship can be plastic due to environmental factors, whether or not male courtship is plastic and depends on social environment, such as the genotype of the female, is relatively understudied (Chaine and Lyon 2008; Wilgers and Hebets 2011). If animals can adjust their mating courtship based on female identity, this might influence how selection leads to divergence in behavior between populations and species.

The fruitfly *Drosophila melanogaster* has been extensively studied for both sexual behavior and social behavior (Greenspan and Ferveur 2000; Auer and Benton 2016), making it an ideal system to investigate what facilitates courtship plasticity. The majority of research conducted in *D. melanogaster* focuses on genotypes that have originated in North American and Europe. Importantly however, genotypes from Southern Africa have substantially divergent mating behavior (Wu, et al. 1995; Ting, et al. 2001; Grillet, et al. 2012; Jin, et al. 2022). This creates both a need and an opportunity to understand how mating behavior and courtship plasticity evolve among populations.

In non-African *D. melanogaster* there are specific cues that have experienced sexual selection and are highly predictive of mating success, with singing being a critical trait. (Greenspan and Ferveur 2000; Ferveur 2010; Pavlou and Goodwin 2013; Yamamoto and Koganezawa 2013). During the initial stages of courtship, males sample pheromones on the female using gustatory receptors on the foretarsi (Lu, et al. 2012; Toda, et al. 2012), with 7,11-heptacosadiene being the key female pheromone that males sense to initiate courtship. This signal is relayed from the foretarsi to the brain where P1 and pIP10 neurons control the singing behavior (Von Philipsborn, et al. 2011). Male singing then induces females to be receptive to copulation (Wang, et al. 2020; Wang, et al. 2021). Male singing is thus a key mating trait and has been extensively studied (Billeter, et al. 2009; Ruta, et al. 2010; Toda, et al. 2012; Pavlou and Goodwin 2013).

When considering Zimbabwe genotypes there are several important differences in courtship and the circuit for singing. Zimbabwe males sing less with the total amount of singing differing between genotypes (Jin, et al. 2022). Zimbabwe males perform other behaviors, such as scissoring and circling (Jin, et al. 2022) that are absent in non-African *D. melanogaster* but seen in some closely related species (Cobb, et al. 1985). The role of these behaviors in *D. melanogaster* is currently unknown.

Female pheromonal differences may be a key driver of these behavioral differences between Zimbabwe and non-African *D. melanogaster* males. Instead of the main female pheromone being 7,11-HD, Zimbabwe females produce an isomer, 5,9-HD, as the most abundant compound (Coyne, et al. 1999; Dallerac, et al. 2000). While Zimbabwe females do produce some 7,11-HD, the quantity is significantly less than non-African genotypes. The gene underlying this female pheromone difference is a desaturase, *desat2*, that changes the position of double bonds in the hydrocarbon molecule. The majority of non-African genotypes have a mutation in *desat2* that creates a null allele with no expression. The lack of expression results in the inability to synthesize 5,9-HD (Coyne, et al. 1999; Dallerac, et al. 2000). Correlated with this female pheromone difference is a difference in female mating behavior.

Zimbabwe females have strong reproductive isolation and reject non-African males, whereas non-African females show no preference for Zimbabwe or non-African males (Wu, et al. 1995; Ting, et al. 2001). Differences in both male pheromones and male singing could play a role in reproductive isolation in this system (Grillet, et al. 2012; Jin, et al. 2022). More recently it has been shown that the time males spend singing is plastic. Zimbabwe males sing very infrequently to Zimbabwe females and increase the time spent singing when courting non-African females with high levels of 7,11-HD. In contrast, non-African males sing less when courting Zimbabwe females with low levels of 711-HD (Jin, et al. 2022). This plasticity could be important in the evolution of asymmetrical reproductive isolation in this system.

In this study we set out to determine if this male behavioral plasticity is a result of the differences in female pheromones between these populations. We test this hypothesis by decoupling the two female Zimbabwe phenotypes of strong choice and the presence of 5,9 HD by creating a *desat2* mutant in an otherwise pure Zimbabwe background. Using this Z53 *desat2* null mutant genotype we can ask specifically about the role of female pheromones contributing to male courtship plasticity. We predicted that if female CHCs are responsible for the changes in male singing behavior then Zimbabwe males would show increased singing to Z53 *desat2* null females compared with wild type Z53 females. If we instead found that Zimbabwe males court wild type Z53 and Z53 *desat2* mutants females using the same behaviors, then we would conclude that males are using cues other than pheromones to make courtship decisions.

## Methods

### Drosophila stocks

All stocks used in this experiment were grown at 25°C with 50% relative humidity (RH) on a 12:12 light:dark cycle. Molasses medium was used for all experiments. The non-African genotype used was DGRP-882 (Dembeck, et al. 2015; Pool 2015). This genotype randomly mates (i.e. no-preference between African and non-African males) and has 7,11-HD as the dominant female pheromone (Jin, et al. 2022). The African genotype that was used was Z53. As a male this genotype shows courtship plasticity (Jin, et al. 2022). On average Z53 males sing less than non-African males and increase singing when exposed to non-African females. Z53 females show strong behavioral preference for African males and the major female CHC is 5,9-HD. This African genotype was the genetic background for our CRISPR mutation of *desat2*.

### Generation of the desat2 null mutant

The gene *desat2* contributes to female pheromone production by changing the position of double bonds (Dallerac, et al. 2000). In non-African *D. melanogaster* there is a naturally occurring mutation in the upstream region that abolishes expression, creating a null mutation (See above, (Dallerac, et al. 2000; Fang, et al. 2002). However, there may have been additional changes to this locus that have evolved in non-African populations. For this reason, we decided to make a null mutation by removing the coding sequence of the gene in the African genotype Z53. We used homology directed repair to replace the coding sequence in this strain with a visible marker. All primers used are listed in Supplemental Table 1. We amplified 1.5kb homology arms directly from the Z53 strain both upstream and downstream of the CRISPR cut site that spanned almost the entire annotated region of *desat2*. We used PCR to amplify a 3XP3-DSRed region from the plasmid pHD-DSRed ((Gratz, et al. 2014); Addgene #51434). These pieces were assembled into a single plasmid using Gibson Assembly. We used Target Finder (Gratz, et al. 2014) to search for two cut sites around *desat2* and designed corresponding sgRNA primers. Oligos to make the sgRNA were cloned into the pucU6 plasmid following the protocol from flycrispr.org. Injections of the donor plasmid, sgRNA, and Cas9 protein were conducted by Genetivision. We verified the insertion and successful transformation by scoring the 3XP3-DsRed marker and by PCR amplification of the insert in transgenic individuals.

### Phenotyping the desat2 mutant

After ensuring correct orientation of the insertion and removal of *desat2* sequence we made a homozygous line. We found no expression of *desat2* using RT-PCR (Supplemental Fig. 1). To quantify the changes in female pheromones we extracted and quantified CHCs as described (Jin, et al. 2022). We also extracted CHCs from DGRP882 and Z53 females for comparison (Fig. 1B). We used a choice test to determine the mating phenotype of *desat2* null females as described (Roy and Castillo 2024). Virgin female *desat2* null females were aged for 7-10 days and paired with a wild type Z53 male and wild-type DGRP882 male. Each trial was observed for 30 minutes. After mating occurred the remaining male was removed and identified. We distinguished males by feeding them food dye which does not affect female preference (Roy and Castillo 2024).

**Figure 1.**
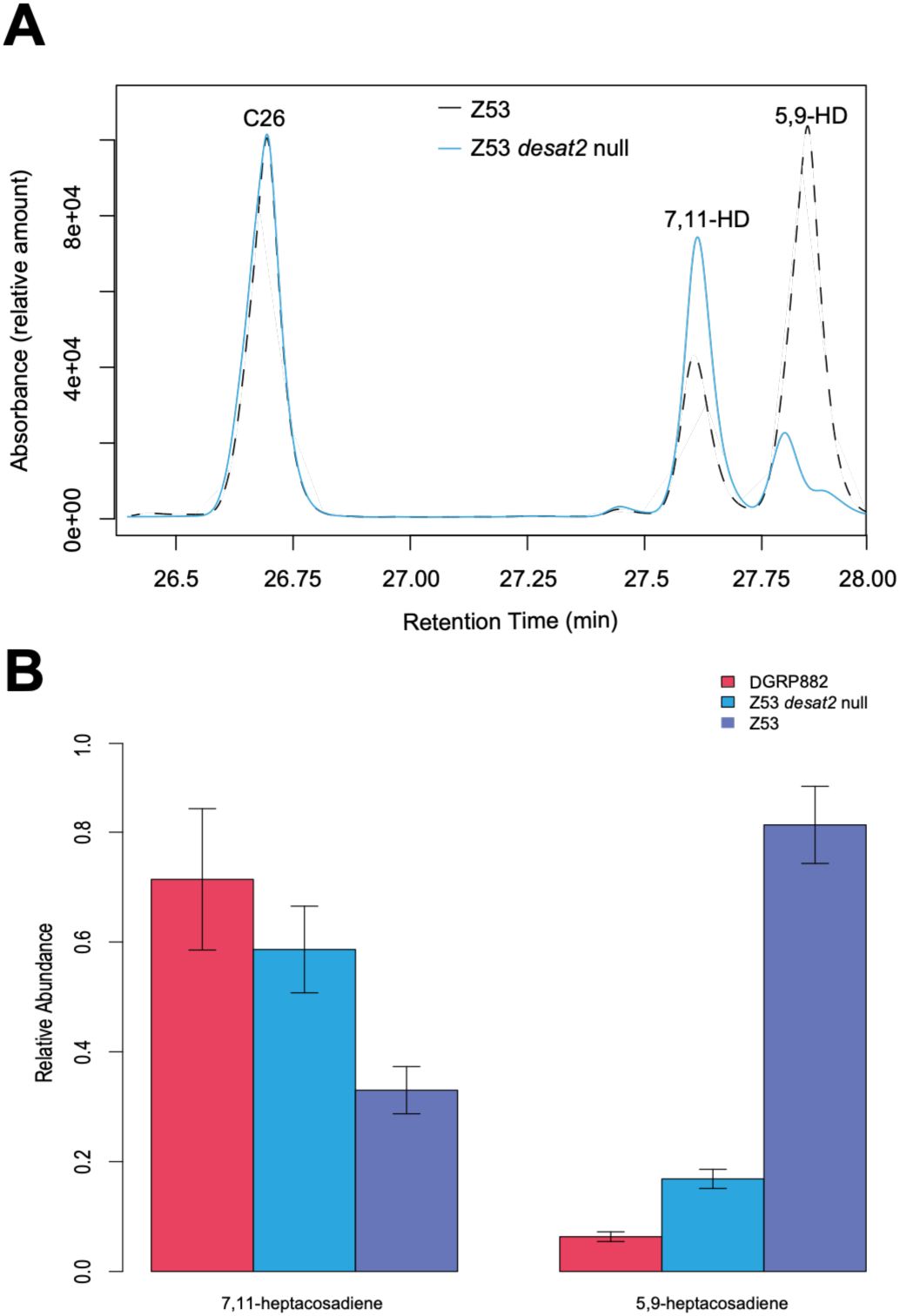
The cuticular hydrocarbon differences between the Z53 *desat2* null mutant and wild-type strains. A) The region of a chromatograph layout focusing on the heptacosadiene (HD) isomers. The Z53 strain produces 5,9-HD as the major compound with some 7,11-HD. 5,9-HD is completely absent in the Z53 *desat2* null mutant, the remaining peak is a co-segregating compound (Coyne et al 1999). B) Quantification of 7,11-HD. 5,9-HD, and co-segregating compounds from replicate females (DGRP-882 n=3, Z53 n=5, Z53 *desat2* null n=4) showing the shift in the Z53 *desat2* null female to resemble the non-African DGRP882 female. All compounds with a retention time between 27.75 and 28 minutes were combined into the 5,9-HD peak following Coyne et al (1999).

**Figure 2.**
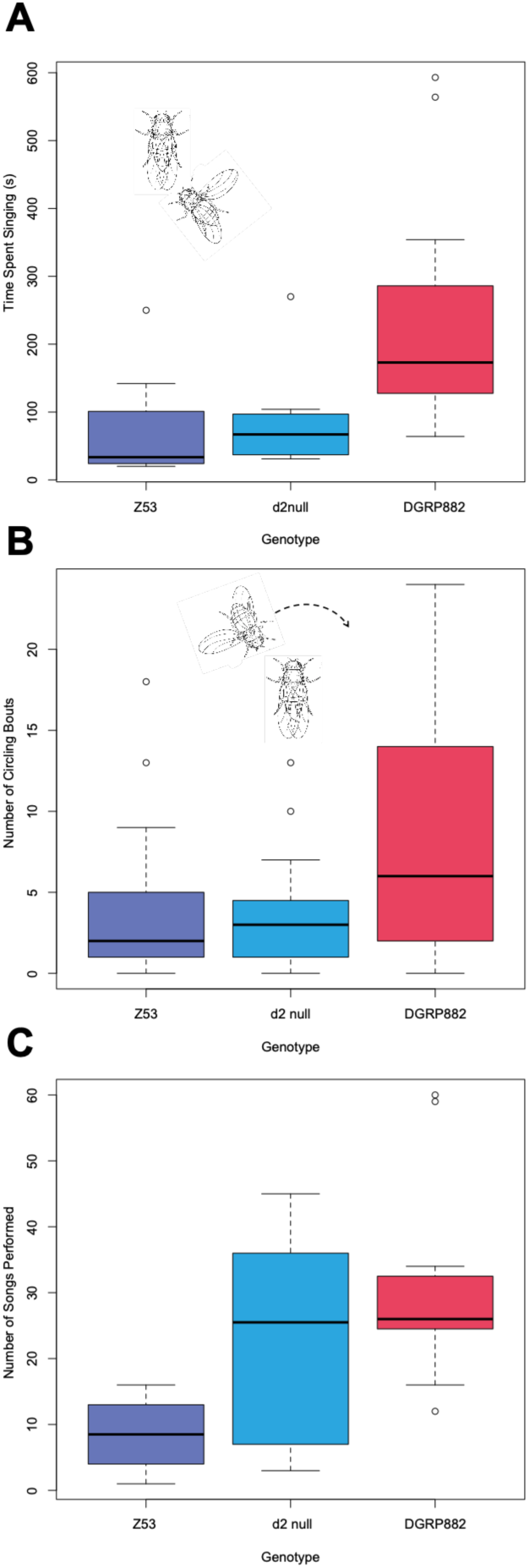
Z53 males treat Z53 *desat2* null females mostly like a wild type Zimbabwe female. A) Z53 males spend the same amount of time singing to Z53 *desat2* null females as Z53 females B) Z53 males perform the same number of circling bouts to Z53 *desat2* null females and Z53 females. The inset in each panel represents the general orientation of the male and female individual during this courtship behavior. C) The number of songs performed to each female genotype showing more songs performed to Z53 *desat2* null females compared to Z53 females. DGRP-882 n=26, Z53 n=21, Z53 *desat2* null n=19.

### Courtship experiments

We assayed courtship plasticity using Z53 males because they actively court and mate with all females. We wanted to avoid issues of males being rejected by females in case this altered their courtship behavior (Balaban-Feld and Valone 2018). Individual males were paired with DGRP882 females, Z53 females, or the *desat2* null females. Individuals were collected as they eclosed as virgins and aged for 7-10 days. Males were placed in vials individually the day before mating (Jin, et al. 2022). All behavioral trials were conducted in a room that had temperature and humidity controlled and matched the incubator (25°C and 55% RH). Temperature and humidity control are important because both are known to effect courtship behaviors including singing (Ritchie and Kyriacou 1996; Noor and Aquadro 1998).

On the day of the mating experiment individuals were added to the courtship chamber without anesthesia. The chamber allows male and female individuals to be isolated until the experiment begins. Pairs were allowed to interact, court, and mate with video recorded for 30 minutes. We used a light box for standardized lighting and we ensured that humidity and temperature in the light box were consistent with the humidity-controlled room.

### Quantification of behavior using ethograms for mated pairs

To determine if courtship behavior of the males was different based on the female genotype we scored behaviors using the BORIS software (Friard and Gamba 2016). The behaviors that we scored were chasing, singing, scissoring, circling, grooming, and attempted copulations (Jin, et al. 2022). Manual scoring was essential for scissoring and grooming behaviors as these are not easily identified by machine learning classifiers (see below). Singing and chasing were both scored as state variables, where the time preforming the behavior is recorded, as well as point variables, where the performance of the behavior is a discrete variable. The remaining behaviors were all scored as point behaviors.

Before scoring for behavior all videos were screened for pairs that successfully copulated. We also selected videos where males and females did not have wing damage or mobility issues. We selected 10 videos per genotype combination. After this filtering each observer watched the video and scored behavior without knowing the genotype of the female. After behavior was scored, we matched the observation with the genotype of the specific pairs observed.

### Quantifying courtship behavior using machine learning classifiers for all replicates

We followed the protocol outlined in DANCE to quantify courtship using behavioral classifiers and machine learning (Yadav, et al. 2025). Our goal was to increase sample size and look at potential differences in singing and courtship plasticity. Using DANCE, videos are first processed by cropping so that only a single well is in view. These cropped videos are used to track individuals with FlyTracker (Eyjolfsdottir, et al. 2014). Finally the behavioral classifiers that were trained through DANCE were used to score attempted copulations, following, circling, and singing in JAABA (Kabra, et al. 2013). We passed all available videos through this workflow, however some failed to be scored due to errors in processing, for example low-quality video where individuals could not be tracked. For this reason, there is a large, but not completely overlapping, set of replicates between the manually and automated scored videos. For the automated analysis we did not pre-filter based on whether courtship ended in successful copulation. This was to increase the number of available observations. We later determined that there was no difference in behavior between pairs that mated or did not mate (see results). There were potential differences in the time spent courting for pairs that did or did not copulate. For this reason, we calculated the time courting as the time the behavioral classifiers observed the first behavior until copulation, or the last time a behavior was observed. We normalized the time-based variables by the time spent courting.

### Statistical analysis

To analyze differences in courtship plasticity for our manually scored data we used linear and generalized linear models that tested the effect of female genotype comparing the replicates that had *deast2* null or DGRP882 females to Z53 females as baseline. For the count data we used a generalized linear model with a Poisson link function and a continuous time covariate. Any differences in the number of behaviors performed over time would result in a significant female genotype effect. The time spent singing was analyzed using a linear model with *desat2* null and DGRP882 genotypes as binary regression coefficients.

To analyze the data from the automated analysis we needed to remove outliers that were assigned spurious behaviors by the classifiers. For example, some videos were scored as having an extreme number of point behaviors like circling. We first used principal components to identify outliers in this multivariate space and removed them. We then looked for outliers for singing and circling behaviors using qqplot in R. This initial filtering was conducted without considering genotype. After this filtering we used the same models described above. We also used a t-test to determine whether there was any effect of copulation status on behavior for each female genotype. All statistical analyses were completed using R v 4.3.3.

### Comparison of behavioral scoring methods

The manual and automated methods of scoring behavior each had advantages and limitations. For traits that were scored using both methods the results were consistent (see below). Traits that were scored by a single method were typically scored manually due to the scale of the behavior and current lack of trained classifiers. The automated behavioral analysis was not able to score small scale behaviors such as grooming, where an individual rubs their foretarsi together, and tapping, where the male touches the foretarsi to the abdomen of the female. These behaviors are difficult for automated analysis given the small size of the *Drosophila* tarsi. In our manual scoring of grooming, we did not see a statistical difference for males courting different female genotypes (Table 1; Supplemental Table 2). Another behavior that was only scored using the manual analysis was scissoring. The DANCE workflow has also not been trained to detect scissoring. While non-African *D. melanogaster* will perform this behavior (McKinney, et al. 2026; Palmateer, et al. 2026), it is more common in African males (Jin, et al. 2022 and closely related species D. simulans {Palmateer, 2026 #133) (Palmateer, et al. 2026). We observed this behavior in our analysis but there were no differences between genotypes (Table 1; Supplemental Table 2).

**Table 1.** The average courtship behaviors for traits scored solely with manual analysis. Z53 males were allowed to court each female genotype for 30 minutes. Standard errors are in parentheses. Chasing is in seconds; all other behaviors are the number they were performed. Bold values are significantly different from the Z53 female baseline using generalized linear regression.

| Courtship Behavior | Z53 | *desat2* null | DGRP882 |
| --- | --- | --- | --- |
| Number of Songs | 8.6 (1.6) | **23.7 (4.5)** | **30.1 (4.6)** |
| Chasing (s) | 40.0 (21.3) | 38.1 (22.6) | 54.27 (25.3) |
| Scissoring | 2.3 (0.76) | 2.5 (0.67) | 3.3 (0.95) |
| Grooming | 4.2 (1.68) | 4.2 (1.42) | 4.6 (1.33) |

The main advantage of the automated analysis was the efficiency of processing large numbers of videos. The automated analysis is also less likely to be affected by observer bias as all videos were scored using the same classifiers. Idiosyncrasies of the classifiers were able to be accounted before in post processing data analysis. For example, the automated analysis tended to overestimate the number of singing attempts as a point behavior. Nevertheless, the estimates for the time spent singing were similar between both analysis methods. This overestimation of singing bouts is likely due to the classifiers assigning one longer singing event as several smaller events. This pattern was also observed with attempted copulations, where what would be scored as one attempt manually would be scored as multiple attempts. In some cases an actual copulation was scored as several attempts. These overestimations were easy to remove with outlier analysis, plotting histograms or qqPlots of these variables and could be removed before comparing these behaviors between genotypes.

The automated classifier was also very accurate in its ability to differentiate singing and circling behaviors. While both behaviors involve wing extensions, they are primarily differentiated by the relative positions of the male and female individuals (Cobb, et al. 1985). The number of circling attempts was similar for the automated analysis and the manual scoring (Table 3). In manual scoring this differentiation can be somewhat subjective, however in the automated analysis this was more precisely defined by using the relative angle between the male and females.

## Results

### desat2 null females have an altered CHC profile but display normal female preference

We quantified cuticular hydrocarbons (CHCs) of our Z53 *desat2* null females to assess the impact of this mutation on CHC production. As expected, the Z53 *desat2* null females resembled the non-African strain for the amounts of 7,11-heptacosadiene (7,11-HD) and 5,9-heptacosadiene (5-9,HD). There was a shift from producing 5,9-HD towards producing 7,11-HD (Figure 1). The amount of 5,11-HD produced by Z53 *desat2* null females was significantly lower than Z53, the progenitor strain (*β*=-0.644, *P*<0.001), and was indistinguishable from the non-African DGRP-882 strain (*t*=-1.289, *P*=0.229). Likewise there was a significant increase of 7,11-HD in Z53 *desat2* null females compared to Z53 females (*β* =0.384, *P*=0.007) with no significant difference compared to DGRP-882 females (*t*=1.09, *P*=0.303).

We found no effect of the Z53 *desat2* null mutation on female mating preference when homozygous. Z53 *desat2* null females displayed strong preference like the progenitor Z53. In choice tests when paired with a Z53 male and a DGRP-882 male, the Z53 *desat2* null females chose Z53 males at a high rate. In 23 replicates, 17 Z52 *desat2* null females chose Z53 males (Binomial test; *P*=1.56e-5) and 6 did not mate. Not mating is a common observation for the Z53 line. As a control, we tested wild type Z53 females and found that in 25 replicates 20 chose Z53 males (Binomial test; *P*=1.97e-6) and 5 did not mate. In no-choice tests over a 24-hour period, Z53 *desat2* null females mated with DGRP882 at a low frequency, 3 out of 20 mated, whereases 20 out of 20 replicates paired with Z53 mated (Fisher exact test; *P*=2.57e-8)

Combined these data demonstrate that the Z53 *desat2* null affects female pheromone profile but not female behavior. We thus can use this genotype to determine how CHC differences influence male courtship.

### African males exhibit plastic courtship behavior

We allowed Zimbabwe Z53 males to court three female genotypes: Zimbabwe Z53 females, non-African DGRP-882 females and the Z53 *desat2* null females. To confirm the presence of courtship plasticity we first examined differences between males courting Z53 and non-African DGR-P882 females. In both the manual and automated analyses the Z53 males sang significantly more to the DGRP-882 females compared to the Z53 females (Fig 1; Table 2; Supplemental Table 2 and 3). The average time spent singing to Z53 females was 74.9 seconds (± 32.8 SE) and the average time singing to DGRP-882 females was 239.0 seconds (±55.9 SE), which were statistically different (β=164.10; *P*=0.0056). Similarly, males performed circling behavior towards DGRP-882 females significantly more compared to Z53 females (β=0.7376; *P*=0.0058). Lastly, males had fewer attempted copulations with DGRP-882 females compared to the Z53 females for the manually scored videos (β=-0.6549; *P*=0.0360). Together these results recapitulate the plasticity that we had previously observed (Jin, et al. 2022). This plasticity did not depend on whether males successfully copulated within the observation time as there was no difference in singing between these replicates for each female genotype. (Supplemental Table 4).

**Table 2.** The average courtship behaviors for traits scored in both the manual and automated analyses. Z53 males were allowed to court each female genotype for 30 minutes. Standard errors are in parentheses. In the manual analysis singing is in seconds. For the Automated analysis this was changed to a proportion of time relative to the total courtship time. All other behaviors are the number of times the behavior was performed. Bold values are significantly different from the Z53 female baseline in generalized linear regression models.

|  | Manual Analysis |  |  | Automated Classifier Analysis |  |  |
| --- | --- | --- | --- | --- | --- | --- |
| Behavior | Z53 | *desat2* null | DGRP882 | Z53 | *desat2* null | DGRP882 |
| Singing | 74.9 (23.8) | 82.5 (22.4) | **239.0 (55.9)** | 0.10 (0.02) | 0.09 (0.02) | **0.19 (0.03)** |
| Chases | 2.4 (0.77) | 1.50 (0.50) | 2.27 (0.93) | 2.26 (0.86) | 0.78 (0.18) | 2.86 (1.14) |
| Circling | 2.0 (0.64) | 2.1 (0.75) | **4.18 (0.94)** | 5.26 (1.69) | 3.81 (1.62) | **9.05 (2.31)** |
| Attempts | 2.80 (1.02) | 1.7 (0.77) | 1.45 (0.85) | 1.39 (0.24) | 1.59 (0.48) | 2.28 (0.76) |

### African males do not use cuticular hydrocarbons in courtship plasticity

We hypothesized that if males were using the cuticular hydrocarbon 7,11-HD as the cue for increased singing to non-African females, then they would increase singing to the Z53 *desat2* null females compared to Z53 females. Z53 males treated the Z53 *desat2* null female like a typical Zimbabwe female. Males actively courted Z53 *desat2* null females, but did not increase their time spent singing (Table 2; Supplemental Table 2 and 3). The time spent singing to Z53 *desat2* null females was 82.5 seconds (± 22.4) and was not significantly different than the time spent singing to Z53 females (*β*=7.60; *P*=0.8931). This was mirrored in circling where there was no significant difference between Z53 *desat2* null females and Z53 females (*β*=0.0488; *P*=0.8759).

The main difference that we observed for males courting Z53 versus Z53 *desat2* null females was in the number of song attempts. In the manual scored analysis there were significantly more song attempts for the Z53 *desat2* null females compared to the Z53 females (β=1.013; *P*=8.1e-16). Males attempted 8.66 songs (±1.8) towards Z53 females and attempted 23.7 (±4.5) songs towards Z53 *desat2* null females. The number of attempts towards Z53 *desat2* null females was similar to the DGR-P882 females (30.1 ± 4.6 song attempts). Since the total time spent singing was not different, this suggests that males initiated and then stopped singing bouts towards Z53 *desat2* females. Males copulated with all three at similar rates: with 42% of the DGRP882 females, 36% of the *desat2* null females, and 38% of the Z53 females within the 30 min observation time.

## Discussion

We used a female genotype that we created to de-couple two important female traits and showed that African male courtship plasticity is not dependent on female pheromones (CHCs). This suggests that male *Drosophila* are receiving other cues from females to adjust their mating behavior. We initially predicted that since Z53 *desat2* null females had a high level of 7,11-HD this would elicit a strong singing response, but this is not what we observed. While Zimbabwe males exhibited courtship plasticity with females from Z53 and DGRP-882 populations, they treated Z53 *desat2* null females like Z53 females. Even though the compound 7,11-HD is paramount for mating in non-African populations it appears to have a different or reduced role in mating interactions in African populations.

The female-specific cuticular hydrocarbon 7,11-HD is thought to be a potent aphrodisiac and an essential part of mating as it elicits male singing (Ruta, et al. 2010; Toda, et al. 2012). This previous work, however, has been completed exclusively in non-African lab strains. African genotypes have different major CHC profiles and different male courtship patterns (Colegrave, et al. 2000; Chertemps, et al. 2006; Grillet, et al. 2012; Jin, et al. 2022). The creation of the Z53 *desat2* null genotype was critical for creating a genotype that breaks the natural correlation between female preference behavior and female CHC that occurs in *D. melanogaster* (Dallerac, et al. 2000; Ting, et al. 2001; Grillet, et al. 2012). The Z53 *desat2* null strain had significantly lower levels of 5,9-heptacosadiene, the dominant Zimbabwe pheromone, and had a significant increase in 7,11-HD. While our study focused primarily on this major female CHC component, other CHCs could still signal Zimbabwe female identity. Our data on the number of song attempts suggest that males sensed the 7,11-HD signal and this did not change their courtship strategy. Zimbabwe males initiated singing to Z53 *desat2* null females more often than to Z53 females, but then stopped singing. The initiation of singing is likely from the 7,11-HD cue, and the stopping of singing could be a behavioral cue. These specific CHC and pheromonal cues are just one component of the complex behavioral dynamics that govern female interactions during courtship.

While most work on female mating has focused on receptivity, such as allowing male access (Grillet, et al. 2006; Wang, et al. 2021), females also perform active behaviors to reject males (Aranha and Vasconcelos 2018). These behaviors might be significant in Zimbabwe mating interactions. In our experiment, males successfully mated with all female genotypes, and we noticed no differences in courtship behavior between males that mated and those that did not. This suggests female receptivity alone did not alter male courtship. Although our video recordings were not optimized to effectively capture or analyze female behaviors, future studies should explicitly investigate these potential differences in female mating behavior. Female mating behavior is often reduced to a binary mated vs non-mated, but females could play an active role in providing males with feedback. This could in turn shape the evolution of reproductive isolation through courtship plasticity.

Differences in plasticity of Zimbabwe and non-African males to 7,11-HD could in part underlie reproductive isolation between these genotypes. Previous results suggest that Zimbabwe females use the time spent singing to reject non-African males (Jin, et al. 2022). Even though non-African males show plasticity in singing, their time spent singing is higher than most Zimbabwe males (Jin, et al. 2022). The non-African males could be strongly stimulated by the 7,11HD compound, consistent with the body of work in lab strains (Billeter, et al. 2009; Ruta, et al. 2010; Toda, et al. 2012; Pavlou and Goodwin 2013), whereas Zimbabwe strains are stimulated but to less of an extent. Differences in sensitivity to other CHCs, such as 7-T, have been recently shown to vary across *D. melanogaster* populations (Ryba, et al. 2026). We did not include non-African males in this experiment because they are rejected by Zimbabwe females and rejection can alter courtship behavior (Balaban-Feld and Valone 2018). Future studies should analyze differences in sensitivity to female CHCs. Whether males sense 5,9-HD or how they perceive it remains unknown, leaving open the possibility that this cue exerts divergent effects on Zimbabwe versus non-African males. The role of plasticity in asymmetric reproductive isolation in this system could suggest a larger role of courtship plasticity in the speciation process.

Asymmetric reproductive isolation is a common pattern driving reproductive isolation between species (Ehrman and Wasserman 1987; Arnold, et al. 1996; Yukilevich 2012). In some cases there has been a loss of a specific cue in one lineage resulting in asymmetrical isolation (Coyne, et al. 1994; Gleason, et al. 2005). However, this mechanism is unlikely to explain the vast majority of cases, especially for closely related species, as premating isolation is often thought to evolve from divergent sexual selection (Panhuis, et al. 2001; Kirkpatrick and Ravigne 2002; Coyne and Orr 2004). In our system, divergent selection has created differences in male courtship yet females still recognize and mate with males of opposite genotypes due to courtship plasticity. Courtship plasticity could be a broadly applicable mechanism explaining patterns of asymmetric isolation across species. When female preference diverges alongside male courtship plasticity, it can lead to asymmetric isolation, depending on the degree of both plasticity and female preference. This evolutionary process is facilitated if male courtship plasticity maintains male fitness across both female genotypes, suggesting either co-occurrence or ongoing gene flow during speciation.

