## Supplemental Tables and Figures for "Female behavioral cues control male courtship plasticity independent of female pheromones in African *Drosophila melanogaster*"

**Supplemental Table 1.** Primers used in the creation of the Z53 *desat2* null mutant. The homology arm primers are based on genomic sequence from the Z53 strain. The left homology arm was designed to amplify 1.5kb of genomic sequence terminating at the g1 CRISPR cut-site. The uppercase sequence is the genomic sequence and lower-case sequence generates overlap with the pHD plasmid that was used as a backbone. The right homology arm starts at the g7 cut-site and amplified 1.5kb. The g7 and g1 cut-sites were chosen based on location within the *desat2* gene and low likelihood of off target cutting. The DSred primers amplified the 3XP3-DSred sequence and had overlap with the homology arms for Gibson Assembly.

| **Primer Name** | **Sequence** | **Description** |
|---|---|---|
| desat2-LA-F | tgtcgcccttcgctgaagcaggtggAGAACGGCATCAAATGGTTCAAC | Forward primer for the left homology arm. |
| desat2-LA-R | tacgataacttcgAGGGGTAGCCGACGC | Reverse primer for the left homology arm |
| DSRed-F | gtcggctacccctCGAAGTTATCGTACGG | Forward Primer for the 3XP3 DSRed |
| DSRed-R | acgtcgggagccaCGAAGTTATACCGGTTAAG | Reverse Primer for the 3XP3 DSRed |
| desat2-RA-F | cggtataacttcgTGGCTCCCGACGTGATC | Forward primer for the Right homology arm |
| desat2-RA-R | cttgaactcgattgacggaagagccTGATCGTGTCCAGCTTTCTG | Reverse primer for the right homology arm |
| g7-F | CTTCGGCCTACGACCTGAAGACGG | Forward primer for the g7 cutsite |
| g7-R | AAACCCGTCTTCAGGTCGTAGGCC | Reverse primer for the g7 cutsite |
| g1-F | CTTCGGATTATATTCCGCCAGACT | Forward primer for the g1 cutsite |
| g1-R | AAACAGTCTGGCGGAATATAATCC | Reverse primer for the g1 cutsite. |

**Supplemental Table 2.** Regression coefficients and P-values from statistical models based on ethograms and manual scoring of behavior  Attempted copulation, circling, following, scissoring, grooming, and the number of songs were modeled as Poisson variables. Singing was a linear model with the response variable being the proportion of time spent singing. Z53 was considered the baseline for these models. Values are bolded if they were significantly different from this baseline.

| Genotype | Behavior | Estimate | Z value | P value |
|---|---|---|---|---|
| Z53 (intercept) | Attempted Copulation | 1.0296 | 5.448 | 5.09e-08 |
| desat 2 | Attempted Copulation | -0.499 | -1.623 | 0.104 |
| **DGRP882** | **Attempted Copulation** | **-0.6549** | **-2.090** | **0.036** |
| Z53 (intercept) | Circling | 0.69315 | 3.100 | 0.00194 |
| desat 2 | Circling | 0.04879 | 0.156 | 0.87591 |
| **DGRP882** | **Circling** | **0.73760** | **2.754** | **0.00589** |
| Z53 (intercept) | Following | 0.87547 | 4.289 | 1.8e-05 |
| desat 2 | Following | -0.47000 | -1.428 | 0.153 |
| DGRP882 | Following | -0.05449 | -0.191 | 0.849 |
| Z53 (intercept) | Scissoring | 0.8329 | 3.994 | 6.48e-05 |
| desat 2 | Scissoring | -0.4274 | -1.288 | 0.198 |
| DGRP882 | Scissoring | 0.3801 | 1.432 | 0.152 |
| Z53 (intercept) | Grooming | 1.435e+00 | 9.301 | <2e-16 |
| desat 2 | Grooming | -4.176e-09 | 0.000 | 1.000 |
| DGRP882 | Grooming | 9.885e-02 | 0.474 | 0.635 |
| Z53 (intercept) | Singing | 74.90 | 1.890 | 0.06915 |
| desat2 null | Singing | 7.60 | 0.136 | 0.89310 |
| **DGRP882** | **Singing** | **164.10** | **2.997** | **0.00566** |
| Z53 (intercept) | Number of Songs | 2.1518 | 19.955 | < 2e-16 |
| **desat 2** | **Number of Songs** | **1.0137** | **8.053** | **8.1e-16** |
| **DGRP882** | **Number of Songs** | **1.2793** | **10.599** | **< 2e-16** |

**Supplemental Table 3.** Regression coefficients and P-values from statistical models looking at behavior that was analyzed from the machine learning and behavioral classifier workflow. Attempted copulation, circling, and following were modeled as Poisson variables with a continuous time co-variate (Frame). Singing was a linear model with the response variable being the proportion of time spent singing. Z53 was considered the baseline for these models. Values are bolded if they were significantly different from this baseline.

| Genotype | Behavior | Estimate | Z value/t-value | P value |
|---|---|---|---|---|
| Z53 (intercept) | Attempted Copulation | -0.7483 | -2.196 | 0.0281 |
| desat 2 | Attempted Copulation | 0.0652 | 0.243 | 0.8077 |
| DGRP882 | Attempted Copulation | 0.0529 | 0.210 | 0.8334 |
| **Frame - Covariate** | **Attempted Copulation** | **2.87e-05** | **3.857** | **0.0001** |
| Z53 (intercept) | Circling | 0.588 | 3.354 | 0.0007 |
| desat 2 | Circling | -0.1385 | -0.835 | 0.403 |
| **DGRP882** | **Circling** | **0.699** | **5.261** | **1.43e-07** |
| **Frame - Covariate** | **Circling** | **2.11e-05** | **5.868** | **4.42e-09** |
| Z53 (intercept) | Following | -1.142 | -3.058 | 0.0022 |
| desat 2 | Following | -0.218 | -0.638 | 0.523 |
| **DGRP882** | **Following** | **0.752** | **2.707** | **0.0067** |
| **Frame - Covariate** | **Following** | **2.918e-05** | **3.842** | **0.000012** |
| Z53 (intercept) | Singing | 0.1359 | 5.249 | 1.91e-06 |
| desat 2 | Singing | -0.0184 | -0.506 | 0.614 |
| **DGRP882** | **Singing** | **0.0993** | **2.948** | **0.004** |

Supplemental Table 4. Comparison of courting behaviors for replicates that did vs did not copulate during the observation time. Each genotype was analyzed separately with copulation status as a predictor variable. The estimate is the regression coefficient for replicates that did copulate. In all behaviors and genotypes there was no affect copulation success on male courtship behavior.

| Genotype | Behavior | Estimate | Z value/t-value | P value |
|---|---|---|---|---|
| Z53 | Attempted Copulation | 0.719 | 0.853 | 0.405 |
| Z53 *desat2* null | Attempted Copulation | -0.286 | -0.824 | 0.422 |
| DGRP-882 | Attempted Copulation | 0.385 | 0.430 | 0.6715 |
| Z53 | Circling | -2.226 | -1.143 | 0.2678 |
| Z53 *desat2* null | Circling | -0.871 | -0.193 | 0.849 |
| DGRP-882 | Circling | 1.321 | 0.434 | 0.668 |
| Z53 | Following | -1.043 | -1.357 | 0.191 |
| Z53 *desat2* null | Following | -1.538 | -0.955 | 0.354 |
| DGRP-882 | Following | -1.489 | -1.348 | 0.191 |
| Z53 | Singing | 0.0045 | 0.119 | 0.906 |
| Z53 *desat2* null | Singing | 0.0311 | 1.057 | 0.305 |
| DGRP-882 | Singing | 0.0539 | 0.856 | 0.400 |

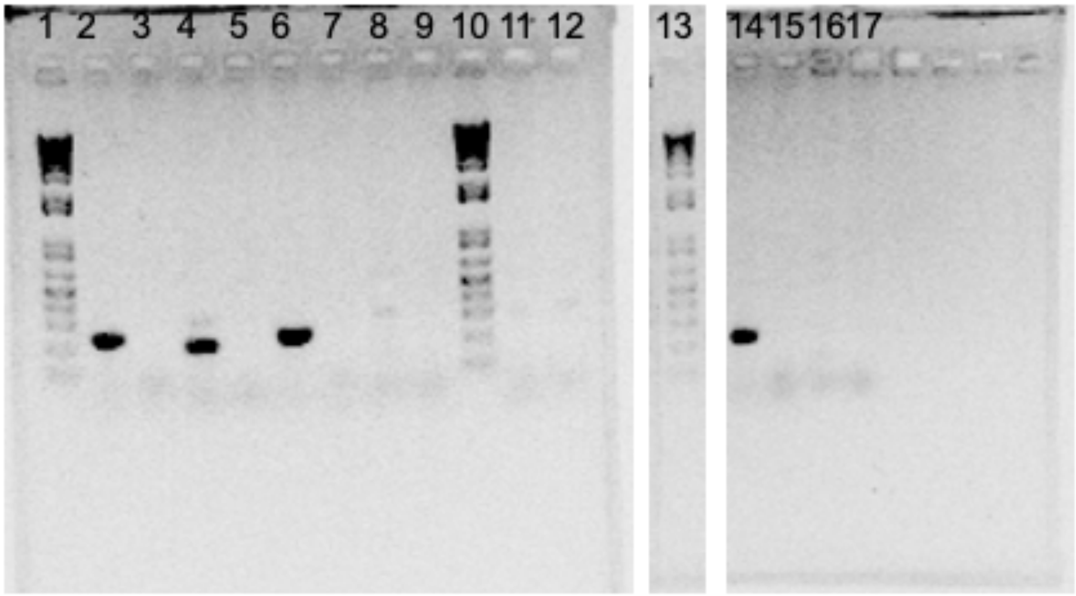

**Supplemental Figure 1.** The Z53 *desat2* null strain lacks *desat2* expression. *desat2* is expressed in Z53 but not DGRP-882 wildtype strains consistent with previous studies. The mutant strain lacks expression which is consistent with the change in cuticular hydrocarbon phenotype. Total DNA was extracted from whole bodies of females and converted to cDNA. For each reaction we also included a RT- control where RNA, buffer, and dT oligos were added but no reverse transcription enzyme. We used the gene *desat1* as a positive PCR control because it is expressed in both genotypes in the oenocyte tissue where cuticular hydrocarbons are synthesized. Z53 is the Zimbabwe genotype and progenitor of the Z53 *desat2* null strain. DGRP-882 is the non-African genotype. The list of samples is as follows.1) 1kb DNA ladder, 2) Z53 female, *desat1* RT+, 3) Z53 female *desat1*, RT- , 4) Z53 female, *desat2* RT+, 5) Z53 female *desat2*, RT- 6) DGRP-882 female, *desat1* RT+, 7) DGRP-882 female *desat1*, RT- , 8) DGRP-882 female, *desat2* RT+, 9) DGRP-882 female *desat2*, RT-, 10) 1kb DNA ladder, 11) d*esat1* PCR neg., 12) *desat2* PCR neg, 13) 1kb DNA ladder, 14) Z53 *desat2* null *desat1* RT+, 15) Z53 *desat2* null *desat1* RT-, 16) Z53 *desat2* null *desat2* RT+, 17) Z53 *desat2* null *desat2* RT-. Lanes 1-12 were run on a single gel. Lanes 13-17 were on a second gel and run using the same PCR master mix as lanes 1-12. There were several lanes between 13 and 14 that were genotypes not used in this paper and removed for clarity of presentation.
